# The TCP transcription factor HvTB2 heterodimerizes with VRS5(HvTB1) and controls spike architecture in barley

**DOI:** 10.1101/2021.04.14.439785

**Authors:** Tatiana de Souza Moraes, Sam W. van Es, Inmaculada Hernández-Pinzón, Gwendolyn K. Kirschner, Froukje van der Wal, Sylvia Rodrigues da Silveira, Jacqueline Busscher-Lange, Gerco C. Angenent, Matthew Moscou, Richard G.H. Immink, G.Wilma van Esse

## Abstract

Barley is the fourth largest cereal crop grown worldwide, and essential for food and feed production. Phenotypically, the barley spike, which is unbranched, occurs in two main architectural shapes: two-rowed or six-rowed. In the 6-rowed cultivars, all three florets of the triple floret meristem develop into seeds while in 2-rowed lines only the central floret forms a seed. *VRS5(HvTB1)*, act as inhibitor of lateral seed outgrowth and *vrs5(hvtb1)* mutants display a six-rowed spike architecture. *VRS5(HvTB1)* is a member of the TCP transcription factor (TF) family, which often form protein-protein interactions with other transcriptional regulators to modulate the expression of their target genes.

Despite the key role of VRS5(HvTB1) in regulating barley plant architecture, there is hardly any knowledge on its molecular mode-of-action. We performed an extensive phylogenetic analysis of the TCP transcription factor family, followed by an *in-vitro* protein-protein interaction study using yeast-two-hybrid. Our analysis shows that VRS5(HvTB1) has a diverse interaction capacity, interacting with class II TCP’s, NF-Y TF, but also chromatin modellers. Further analysis of the interaction capacity of VRS5(HvTB1) with other TCP TFs shows that VRS5(HvTB1) preferably interacts with other class II TCP TFs within the TB1 clade. One of these interactors, encoded by *HvTB2*, shows a similar expression pattern when compared to *VRS5(HvTB1)*. Haplotype analysis of *HvTB2* suggest that this gene is highly conserved and shows hardly any variation in cultivars or wild barley. Induced mutations in *HvTB2* trough CRISPR-CAS9 mutagenesis in cv. Golden Promise resulted in barley plants that lost their characteristic unbranched spike architecture. *hvtb2* mutants exhibited branches arising at the main spike, suggesting that, similar to *VRS5(HvTB1), HvTB2* act as inhibitor of branching. Taken together, our protein-protein interaction studies of VRS5(HvTB1) resulted in the identification of *HvTB2*, another key regulator of spike architecture in barley. Understanding the molecular network, including protein-protein interactions, of key regulators of plant architecture such as VRS5(HvTB1) provide new routes towards the identification of other key regulators of plant architecture in barley.

**Author summary:** Transcriptional regulation is one of the basic molecular processes that drives plant growth and development. The key TCP transcriptional regulator TEOSINTE BRANCHED 1 (TB1) is one of these key regulators that has been targeted during domestication of several crops for its role as modulator of branching. Also in barley, a key cereal crop, HvTB1 (also referred to as VRS5), inhibits the outgrowth or side shoots, or tillers, and seeds. Despite its key role in barley development, there is hardly any knowledge on the molecular network that is utilized by VRS5(HvTB1). Transcriptional regulators form homo- and heterodimers to regulate the expression of their downstream targets. Here, we performed an extensive phylogenetic analysis of TCP transcription factors (TFs) in barley, followed by protein-protein interaction studies of VRS5(HvTB1). Our analysis indicates, that VRS5(HvTB1) has a diverse capacity of interacting with class II TCPs, NF-Y TF, but also chromatin modellers. Induced mutagenesis trough CRISPR-CAS mutagenesis of one of the putative VRS5(HvTB1) interactors, HvTB2, resulted in barley plants with branched spikes. This shows that insight into the VRS5(HvTB1) interactome, followed by detailed functional analysis of potential interactors is essential to truly understand how TCPs modulate plant architecture. The study presented here provides a first step to underpin the protein-protein interactome of VRS5(HvTB1) and identify other, yet unknown, key regulators of barley plant architecture.

## Introduction

Plant architecture is a major determinant for yield and as such has been a target during domestication and breeding. In maize (*Zea mays*) the gene *TEOSINTE BRANCHED1 (TB1)* has been selected during domestication for its role in shaping plant architecture. TB1 inhibits the outgrowth of lateral branches and increased expression of *TB1* in maize resulted in a drastic reduction in number of branches and increased crop yield(1,2). To date, *TB1* orthologs have been targeted for its effect on improved yield in several crops including, pea, potato, barley, rice and wheat(3–7). *TB1* is a member of the plant specific TCP transcription factor family. The family name refers to the founding members *TB1* in maize, *CYCLOIDEA (CYC)*, which is involved in controlling floral bilateral symmetry in snapdragon, and *PROLIFERATING CELL FACTORS (PCF1 &2)* in rice(8,9). PCFs bind to the promoter of *PROLIFERATING CELL NUCLEAR ANTIGEN (PCNA)* to control cell cycle in meristems, as well as DNA synthesis and repair(8). This class of TF exhibits a highly conserved TCP domain, which contains a basic-Helix-Loop-Helix (bHLH) structure involved in DNA binding and protein-protein interactions(8,10). The TCP transcription factor family can be divided into two major phylogenetic clades, class I and class II. TCPs play crucial roles in controlling plant architecture(11,12). In cucumber, for example, mutations in the TB1-clade TCP protein TEN, which contains a highly conserved amino acid sequence only found in Cucurbitaceae, resulted in plants that developed shoots instead of tendrils(13). The maize TB1-clade gene *BRANCHED ANGLE DEFECTIVE 1 (BAD1)* is required for normal tassel branch angle formation(14). The closely related gene in rice, known as *OsTB2* and as *RETARDED PALEA 1, REP1*) controls palea development and floral zygomorphy. TB1, which acts as inhibitor of axillary meristem outgrowth(15– 19), appears to be the most conserved member within the TCP TF family. In barley, *VRS5(HvTB1)* is a key regulator of plant architecture and yield. *VRS5(HvTB1)* is closely related to the maize domestication gene *TB1*. Barley seeds are formed on the inflorescence, which contains the grain producing florets that are arranged on a single main stem, the rachis(20). The rachis develops specialized branches called spikelets, which eventually develop into seeds located on opposite sides of the rachis. Modifications to the overall spike architecture have been vital for cereal domestication and yield improvement(21,22). In barley, the main spike (inflorescence) also underwent significant changes in architecture. For example, wild barley shatters the seeds from the main spike, a characteristic that was lost during domestication of barley(23). To date, the barley spike occurs in two main architectural shapes: two-rowed or six-rowed. In two-rowed lines, only the central floret develops into a seed, in contrast to six-rowed cultivars in which all three florets develop into seeds. *VRS5(HvTB1)* act as inhibitor of lateral seed formation, and as such *VRS5(HvTB1)* has been selected in six-rowed barley cultivars for its role in shaping spike architecture(5). Detailed phenotypical analysis showed that *vrs5(hvtb1)* mutants also exhibit an increased tiller number at early developmental stages(5,24,25). This suggests that, similar to its maize counterpart VRS5(HvTB1) inhibits the outgrowth of lateral branches.

Despite the key roles of VRS5(HvTB1) in barley development, there is hardly any knowledge on the molecular network in which VRS5(HvTB1) is active. Here we performed a comprehensive analysis of barley TCP genes and their chromosomal location. In total, we identified 21 barley TCPs: 11 class I and 10 class II. Given the key roles of TB1 in plant development, we focused on VRS5(HvTB1) and performed an unbiased Y2H screen to identify potential protein-protein interactors and to shed light on its molecular mode of action. This analysis was followed by a more detailed analysis of candidate interactors within the class II TCP TF family. We generated a CRISPR-CAS9 induced mutation in one of the genes encoding a putative VRS5(HvTB1) interactors, *HvTB2*. Our data shows that barley plants that do not have functional HvTB2 develop spikes that lost their characteristic determinate growth pattern. Taken together, our analysis shows that VRS5(HvTB1) has the capacity to heterodimerize with other transcriptional regulators, including closely related class II TCPs. Phenotypical analysis of one of the putative interactors shows that other class II TCPs, besides VRS5(HvTB1), are involved in controlling spike architecture in barley.

## Results

### Barley class II TCPs have a grass-specific sister clade of TB1

To elucidate the phylogenetic relations of the barley TCPs, a maximum likelihood (ML) phylogenetic tree was built including all known members of the barley, wheat, Arabidopsis, rice and maize TCP transcription factor families. With exception of wheat and barley, all sequences were extracted from the iTak(26) and grassius database(27). Wheat TCP genes were extract from Zhao *et al*. (28). For barley and wheat, the TCPs were compared to the newest reference genomes available(29). The multiple sequence alignment was manually curated and non-aligning sequences were removed. For barley, this included HORVU6Hr1G093970.1 which is truncated and not present in the newest reference genome(29) (S1 Table). For wheat, this included TaPCF7.A, TaPCF7.B, TaPCF7.D, which did not contain a TCP domain; and TaTCP19, which was partially truncated. In total 21 barley TCPs, 22 rice TCPs, 24 Arabidopsis TCPs, 62 wheat TCPs and 46 maize TCPs were included in the analysis (S2 Table). Similar to the situation in other plants, barley TCPs grouped into two main classes, class I (PCF) and class II (CIN/CYC/TB1) (Fig 1). Out of the 21 barley TCPs, 19 exhibit a similar genomic organization with either wheat or rice (S1 Fig). Barley TCPs have a close phylogenetic relationship to hexaploid wheat, which contains three copies of each TCP on the A, B and D genomes. Similar to wheat(28), the TB1 locus is duplicated in barley, with a copy on chromosomes 4 and 5 (Fig 1A, S1 Table). VRS5(HvTB1) and TaTB1, both located on chromosome 4, are known regulators of inflorescence architecture. However no function has been attributed to their paralogs, HvTB1-like and TaTB1.2, on chromosome 5.

**Fig 1.**
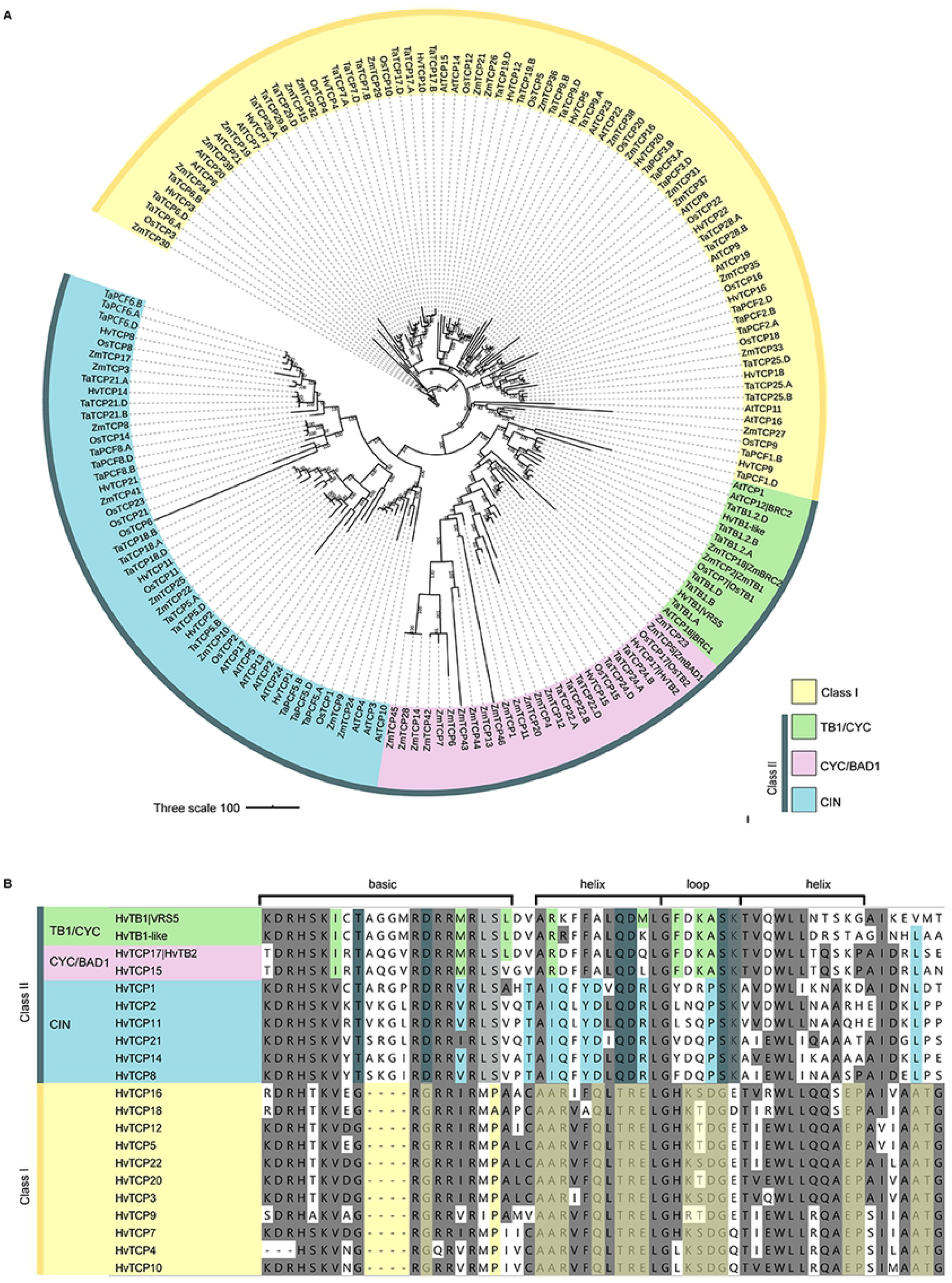
Phylogenetic relationships and sequence conservation of barley TCP transcription factors. (A) Maximum likelihood phylogenetic tree of TCP transcription factors from barley, wheat, rice, maize and Arabidopsis. (B) Amino acid sequence alignments of the TCP domain of barley TCPs. A gray background indicates a high similarity in the conserved TCP domain, independent of class I or II; green blue and yellow background indicates conserved amino acids corresponding to the TB1/CYC, CIN and class I clade, respectively. Purple indicates the two TCPs HvTB2 and HvTB15, which are most closely related to BAD1 according to the phylogenetic analysis.

Barley HvTB2 and HvTCP15 fall together with ZmBAD1 and OsTB2 into a sister clade of TB1 (Fig 1A). To further elucidate the origin of this subclade, we performed a phylogenetic analysis comparing HvTB-like genes in 19 monocot and eudicot plant species. This analysis shows that both HvTB2 and HvTCP15 fall into a grass-specific sister clade of TB1 (S2 Fig). Within this clade, HvTB1 is more similar to ZmBAD1 and to OsTB2, while HvTB15 is most similar to OsTCP15 and the sorghum mutliseeded1 (msd1) TF, which is well known for regulating inflorescence architecture(30). Taken together, similar to maize and rice, the barley and wheat TCP TF families have a grass-specific sister clade.

### VRS5(HvTB1) forms heterodimers with closely related class II TCP T

TCP transcription factors can form homo- and heterodimers, which affect their DNA binding capacity and specificity. To evaluate the protein-protein interaction capacity of barley VRS5(HvTB1) we performed unbiased and targeted yeast two-hybrid (Y2H)-based screenings using this TCP protein as bait. Because of autoactivation of yeast reporter genes, the N-terminal part of the HvTB1 protein was removed (VRS5(HvTB1^NtDEL83^)). Subsequently, we generated a Y2H cDNA expression library of the early and late developmental stages of the barley shoot apical meristem (SAM), respectively (S3 Fig). These stages were selected as VRS5(HvTB1) is highly expressed in the developing SAM (Fig 2B). Screening of VRS5(HvTB1^NtDEL83^) against the barley cDNA libraries resulted in the identification of 114 positive colonies, from which 16 encoded unique proteins in frame with the GAL4 AD-domain (S3 Table). Amongst these are SWItch/Sucrose Non-Fermentable (SWI/SNF) complex subunits, Nuclear transcription factor Y (NF-Y) and HvTCP2.

**Fig 2:**
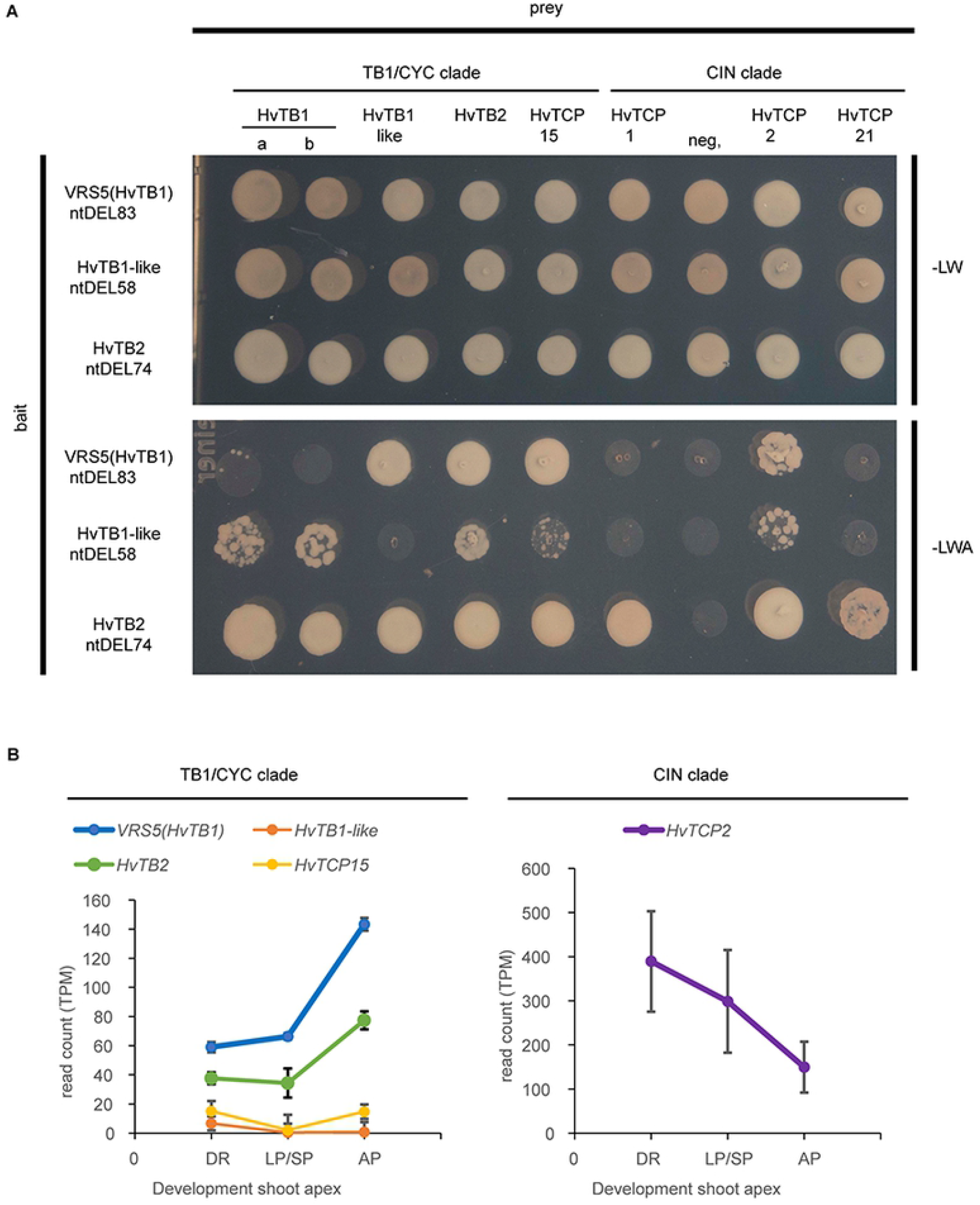
Protein-protein interaction and gene expression of barley TCP transcription factors in the TB1-clade and CIN-clade. (A) Protein-protein interactions were scored on the medium lacking leucine (L), tryptophan (W) and adenine (A), medium lacking L and W was used as positive control for the mating. As negative control (neg.) a barley gene annotated as TF with unknown function was used as prey. For the bait vector, N-terminal deletion constructs of VRS5(HvTB1), HvTB1-like and HvTB2 were used. (B) Expression of TCPs that interact with VRS5(HvTB1) based on transcript per million (TPM). For each RNA-Seq library three independent replicates extracted from GSE102191(33) and GSE149110(34) were re-analysed. DR= double ridge stage; LP/SP is the lemma and stamen primordia stage; AP is the awn primordium stage.

It is well known that TCP proteins interact amongst each other with a preference for interaction with other members within the same clade(31). However, interactions can be easily missed in a library screening. Therefore, we decided to evaluated the interaction between VRS5(HvTB1^NtDEL83^) in the BD vector and against a Arabidopsis TF Y2H library(32) in a heterologous Y2H screen. In this screen, we identified AtTCP1 (AT1G67260) and AtBRC1(AT3G18550), both class II TCPs, as interactors. No interaction was observed with any of the class I TCP proteins, as expected. Within the heterologous screen, we also observed an interaction with AtNF-Y proteins, which confirms the interaction found with barley NF-Y factors in the barley cDNA library screen. Moreover, VRS5(HvTB1) was capable of interacting with Arabidopsis HOMEODOMAIN-containing proteins, and MYB-like transcriptional regulators (S3 Table).

To further evaluate protein interactions of VRS5(HvTB1), we investigated its capacity to form complexes with other class II TCPs in barley. For this we selected as preys: HvTCP1, HvTCP21 and HvTCP2 which belong to the CIN clade; and the four class II TCPs within the TB1 and CYC/BAD1 clade (Fig. 1). In this targeted analysis, two VRS5(HvTB1) protein variants were included, encoded by two natural alleles, the a and b allele, which correspond to the six-rowed and two-rowed cultivars, respectively(5). Because of autoactivation by the selected class II TCP proteins, no complete pair-wise matrix-based screen could be performed. For this reason, we generated N-terminal deletion variants for VRS5(HvTB1), HvTB1-like and HvTB2 and used these truncated proteins as baits. VRS5(HvTB1) and HvTB1-like showed a weak homo- and heterodimerization capacity (Fig 2A, S4 Fig). No difference in this homo- and hetero dimerization capacity was observed between the a- and b allele variants of VRS5(HvTB1). Both VRS5/TB1 and TB1-like proteins showed a consistent interaction with HvTB2, HvTCP15 and HvTCP2 (Fig 2A). Vice versa, HvTB2 interacted with both VRS5(HvTB1) and HvTB1-like. No interaction was observed between VRS5(HvTB1) or HvTB1-like and the CIN-clade proteins, HvTCP1 and HvTCP21. Altogether these experiments revealed that the barley TB1-like TCPs preferentially interact with closely related members within the class II clade of TCP proteins.

For biological relevance, genes encoding interacting proteins should be co-expressed and therefore, we compared the expression patterns and levels of *VRS5(HvTB1)* and of the genes encoding the interacting TCP TF in the developing shoot apex by re-analysing available RNA-Seq data of cv. Bowman apical meristems(33,34). *VRS5(HvTB1)* and *HvTB2* have a low expression at the double ridge stage, which increases in the lemma and stamen primordia stages (LP/SP) and up to the awn primordia stage (AP) (Fig 2B). In comparison, *HvTB1-like* and *HvTCP15* are lowly expressed within the developing shoot apex at all three investigated developmental stages. *HvTCP2* is highly expressed in the shoot apex, but follows an opposite trend in time when compared to *VRS5(HvTB1)* and *HvTB2*. Taken together, HvTB2 follows a similar expression pattern as it’s interaction partner VRS5(HvTB1).

### Barley HvTB2 controls spike branching

HvTB2 is a putative interactor of VRS5(HvTB1), and follows a similar expression pattern when compared to VRS5(HvTB1) (Fig 2). Moreover, our phylogenetic analysis showed that HvTB2 is closely related to maize ZmBAD1 and OsTB2, with similar domain architecture when compared to ZmBAD1 (Fig 1). These observations prompted us to study the function of HvTB2 in more depth and led to the hypothesis that HvTB2 influences inflorescence architecture in barley, at least partially in concert with VRS5(HvTB1). To test this hypothesis, we generated targeted mutations in *HvTB2* using CRISPR-CAS9 gene editing in barley cv. Golden Promise (GP). Aiming at larger deletions and a specific null mutant for this TCP gene, three guides were used, all targeting the N-terminal part of *HvTB2* before the conserved TCP domain (Fig 3A, S5 Fig). In total, 38 CAS9 positive plants were generated, from which one showed a putative biallelic event. Screening of the T2 transformants of this line resulted in two novel *HvTB2* alleles, *hvtb2-1* and *hvtb2-2*, containing a 56bp deletion and a 184bp insertion, respectively (Fig 3A). In both *hvtb2-1* and *hvtb2-2* the mutational event caused a frame shift before the TCP domain, thereby generating full null mutants of *HvTB2*. Both mutants exhibited spikes that lost the characteristic determinant growth pattern, with branches forming on the main rachis (Fig 3B). The seed bearing branches were significantly shorter when compared to the main spike (Fig 3C, Table S5). The outgrowth of branches from the main rachis occurred mainly on the basal part of the spike (S6 Fig). In addition, we also observed that some of the basal seeds in *hvtb2* showed two awns and/or fused seeds, a phenotype that does not occur in the wild type GP (Fig 3B). Due to the reduced spike length, the total number of grains was only moderately increased in the *hvtb2* lines, despite the presence of lateral branches (S7C Fig). The thousand grain weight (TGW) and grain width was significantly reduced in the *hvtb2*-lines (Fig 3D; S7D-S7E Fig). Overall, the grain size in the lateral branches was reduced when compared to the main spike (S7D-S7E Fig). Interestingly, *hvtb2* mutants displayed a significant increase in tiller number when compared to the wild type GP (Fig 3E). Taken together, our data suggest that *HvTB2* influences multiple yield-related traits throughout barley development, similar to VRS5(HvTB1). The macroscopic phenotype resembles previously described phenotype for *intermedium-h* (*int-h*) and *compositum 1* (*com1*)(35,36). Targeted PCR amplification showed no amplicon in the coding sequence or promoter region of *int-h42, int-h*.*43* and *int-h*.*44, int-h*.*83, com1*.*a* and *com1*.*b* (S8A Fig). Two of the lines tested, *int-h*.*83* and *com1*.*c* contained a nonsynonymous mutation that resulted in an amino acid change within the conserved TCP domain (S8B-S8C Fig). Therefore, *HvTB2* is a good candidate gene for the *int-h* and *com1* loci. Taken together, we identified *HvTB2* as gene controlling spike architecture.

**Fig 3.**
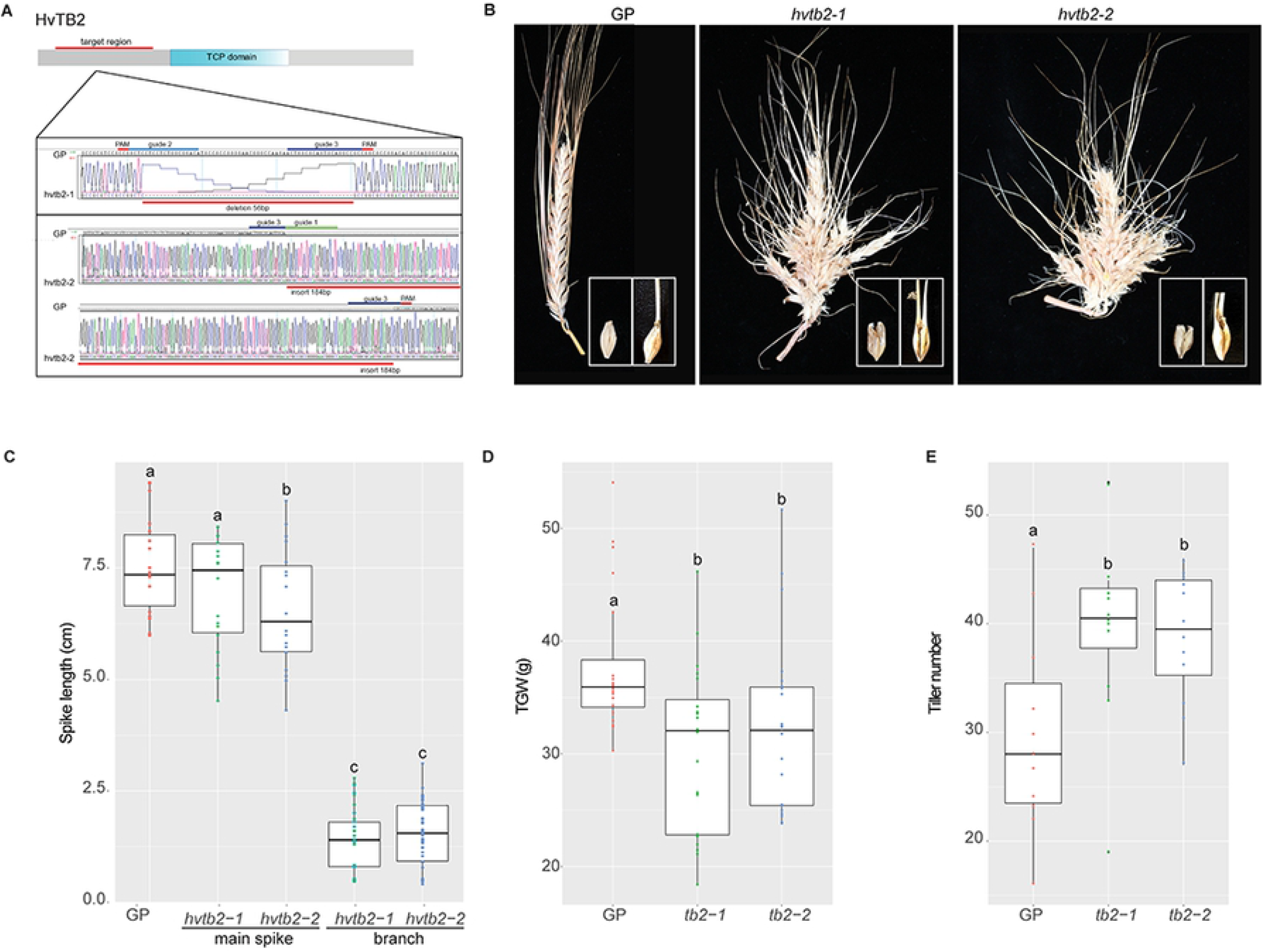
Macroscopic phenotype of HvTB2 mutants. (A) CRISPR-CAS9 target site and *hvtb2* mutants generated. Trace files show the sequence of cv. Golden Promise (GP) in comparison the 57bp deletion mutant, *hvtb2-1* and the 184 bp insertion mutant *hvtb2-2*. (B) Spike phenotype of the wild-type GP in comparison to the generated *hvtb2-1* and *hvtb2-2* mutants. Right corner inset shows an enlarged image of the seeds, with clear split of the awn and fused seeds which is observed in both mutants. (C-E) Spike length, thousand grain weight (TGW) and tiller number measurements of GP, *hvtb2-1* and *hvtb2-2*. Per genotype: spike length n=18 spikes; for tiller number n= 12 plants. TGW is based on extrapolation of the weight of 15 seeds, n= 20 pools. Different letters indicate experimental groups that were significantly based on a one-way ANOVA (P ≤ 0.05), same letters indicate not significant under this criterium.

### *hvtb2* acts as a boundary gene

To determine the origin of the lateral branches that appear on the main rachis we compared the morphology of wildtype GP and *hvtb2* mutants at LP/SP using scanning electron microscopy. In GP the triple spikelet meristem is formed and outgrowth of lateral branches is supressed. In contrast, the central spikelet at the base of the meristem of *hvtb2* mutants is enlarged, resembling a branch meristem instead of a triple spikelet meristem (Fig 4A). This altered development mainly occurs at the basal part of the spike. In line with this the branches in the mature spike are only observed in the first five rachis nodes (S6 Fig). Overall, no major differences were observed in leaf number or the overall developmental speed of the apex (S9 Fig), suggesting that HvTB2 mainly acts on inhibition of the spike branching. The lateral spike branch showed an indeterminate growth pattern, and continued to grow and differentiate after producing the floret meristems. The branch meristem-like structures are still vegetative at the stamen primordium stage (Fig 4A), and start to initiate spikelet primordia when the inflorescence transitions to the awn primordium stage (S9 Fig). No major phenotypes were observed at the double ridge stage (S9A Fig). In line with this, expression of *HvTB2* is low in this tissue and not yet localized to the spikelet primordia (Fig 2, S9 Fig). RNA *In-situ* hybridization shows that, at the awn primordium stage, *HvTB2* mRNA is mainly expressed at spikelet meristem boundaries (Fig 4B, S10 Fig). This suggest that HvTB2 may act within the triple floret meristem as boundary gene.

**Fig 4.**
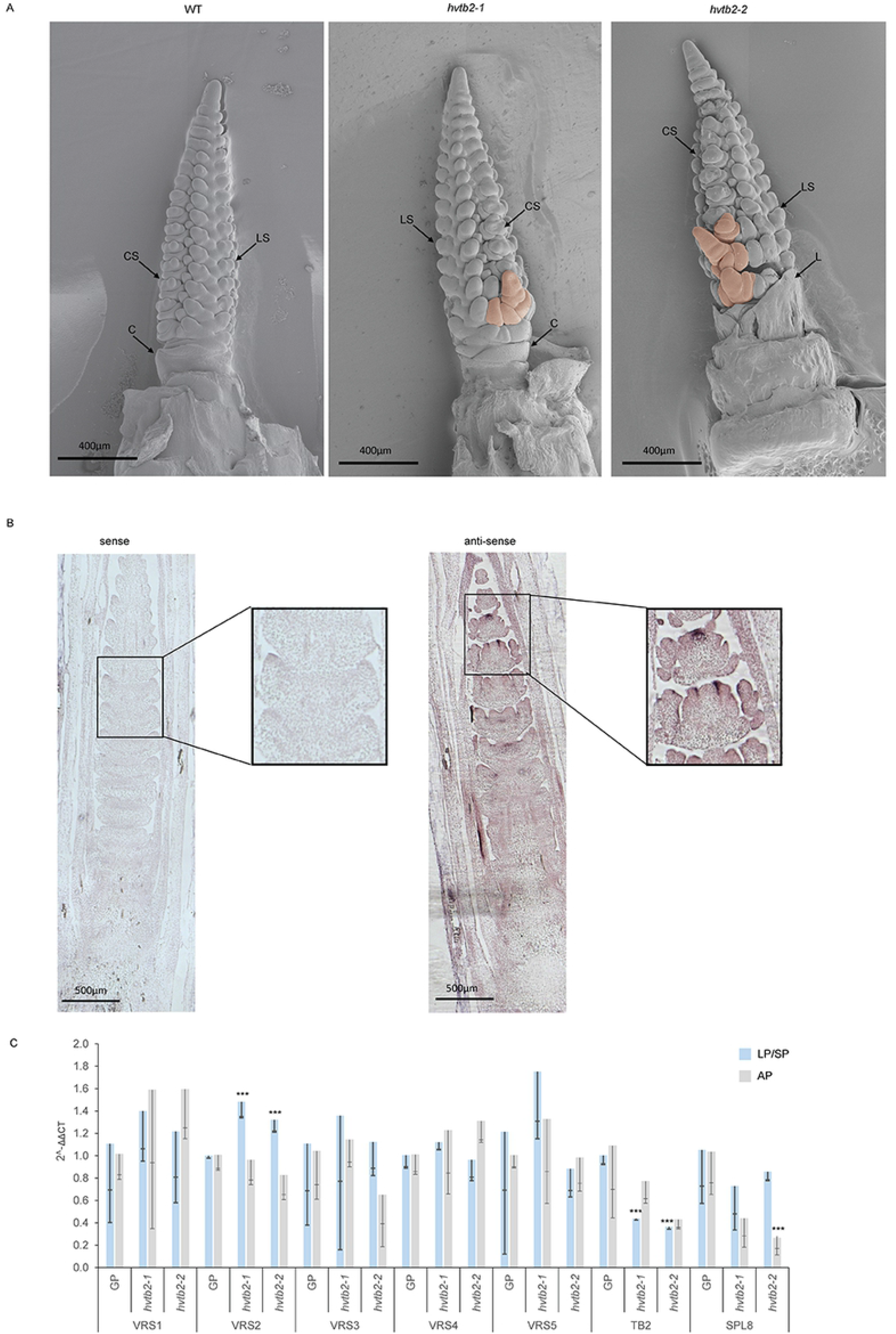
Meristem phenotype of *hvtb2* mutants. (A) Scanning electron microscope images taken at the lemma and stamen primordia stages (LP/SP). Pink color indicates the outgrowing branch structure in the developing meristem. CS = central spikelet meristem; LS= lateral spikelet meristem; C= colar; L=leaf. (B) RNA *in-situ* hybridization of *HvTB2* in the background of cvBowman. Squares in panel are enhanced images in the defined region. (C) RT-PCR analysis of the VRS genes, TB2 and SPL8 in *hvtb2-1* and *hvtb2-2* compared to the wild type cv Golden Promise (GP) at the LP/SP and AP. Statistical differences between the ΔΔCT values was calculated using a t-test using a p-value of 0.05 as the threshold. Asterisks indicate significant differences when compared to GP. For all datapoints n≥3 biological replicates.

Next, we evaluated to what extent HvTB2 influences the expression of other, known regulators of row-type architecture. To this end, we performed RT-PCR analysis in immature shoot apexes of *tb2-1* and *tb2-2* mutants compared to wild type GP lines. Two developmental stages were selected, the lemma and stamen primordia stage (LP/SP) and the awn primordium stage (AP), where at the LP/SP a significant downregulation of *HvTB2* was observed. With exception of *VRS2*, which was significantly upregulated at the LS/SP stage, none of the other row-type genes was changed in expression in neither *tb2-1* nor *tb2-2*. We also included *SQUAMOSA PROMOTER-BINDING-LIKE8* (*SPL8;* HORVU0Hr1G039150)(37). In maize the SPL8-like gene *LIGULELESS 1* (*LG1*), act downstream of ZmRAMOSA2 (RA2) and ZmBAD1(14). Interestingly, SPL8 is significantly downregulated in the *hvtb2-2* mutant at the AP stage, suggesting that like in maize SPL8-like genes act downstream of *hvtb2*. Taken together, our detailed phenotypical analysis indicates that *HvTB2* controls spike determinacy and acts as a boundary gene.

### Barley *HvTB2* is highly conserved

*TB1* is a well-known gene targeted during domestication of several crops including maize, wheat, rice and barley. To evaluate if *HvTB2* is also subjected to selection we performed a haplotype analysis based on available single-nucleotide polymorphism (SNP) (38,39). To assess both natural variation and possible selection through breeding, sequences from cultivars and landraces were included. For comparison, *VRS5(HvTB1)* was also included in the analysis. Our analysis indicates that there are 4 major *VRS5(HvTB1)* haplotypes. Two major haplotypes, *HvTB1*.*a* and *HvTB1*.*b*, are primarily found in 6-rowed and 2-rowed cultivars respectively (S10 Fig), corroborating previous reports(5). Based on the PROVEAN score for conservation analysis no major functional changes are expected by the difference between the *HvTB1*.*a* and *HvTB1*.*b* alleles (S11 Fig). Haplotype analysis on *HvTB2* shows two major haplotypes (HAP1 and HAP2), and six minor haplotypes. Form these, four minor haplotypes did not cause a change in the amino acid sequence when compared to HAP1 (Fig 5). For the other remaining haplotypes, no changes were observed in the conserved TCP domain. Based in the PROVEAN score for conservation analysis no functional changes are expected between the haplotypes (S8C Fig). None of the haplotypes identified were specific for either 2-rowed or 6-rowed cultivars nor for wild barley, landraces or cultivars (Fig 5). Taken together, our results indicates that there is very little variation within the HvTB2 gene.

**Fig. 5.**
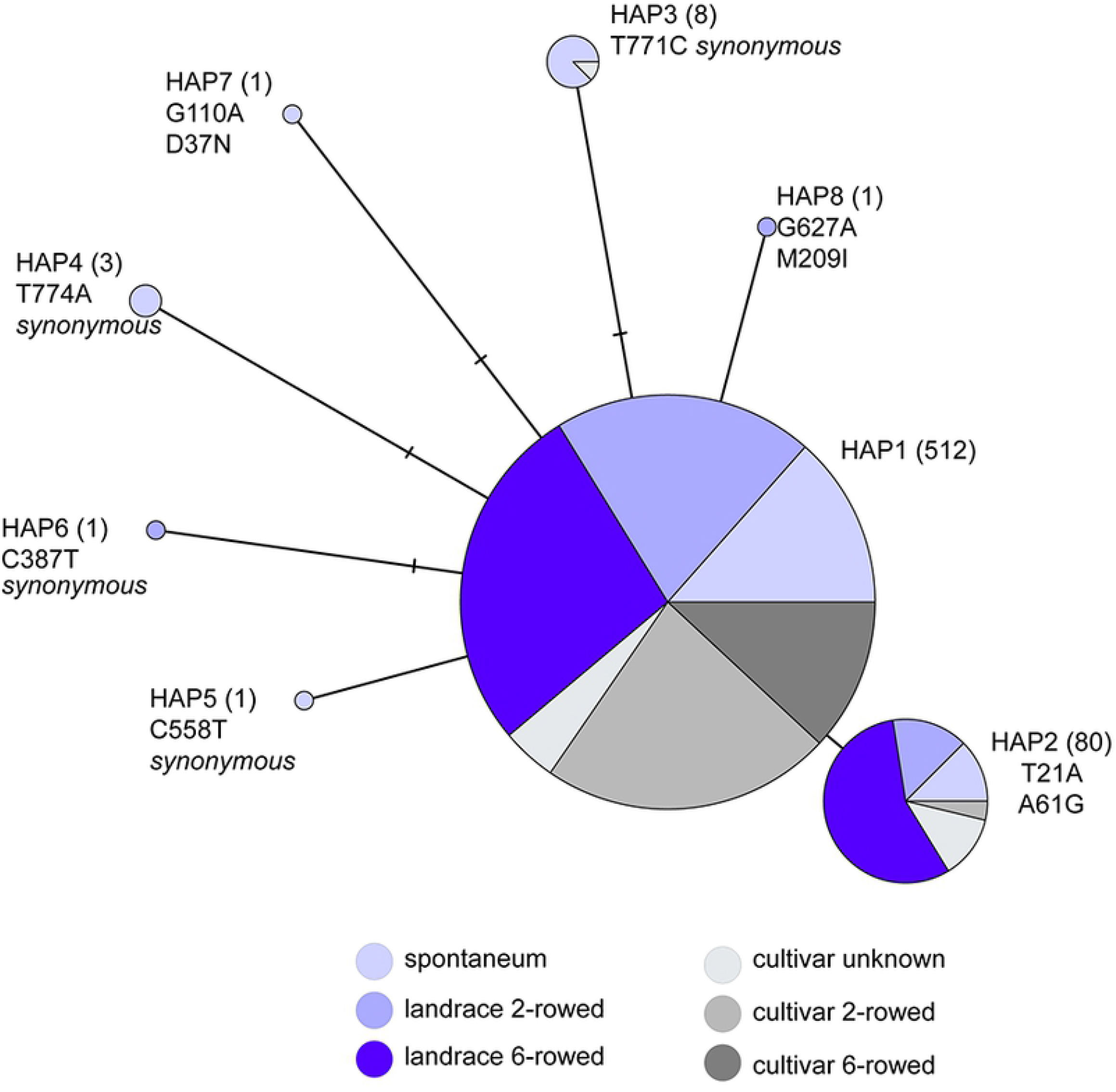
Haplotype analysis of *HvTB2*. Haplotype analysis is done on SNPs present in 607 individual plant lines ranging from wild barley (spontaneum), landraces and cultivars with 2-rowed or six-rowed spike architecture. Number of plants per haplotype is indicated between brackets, SNPs identified and changes that occur at the amino acid level are stated below the haplotype.

## Discussion

TCP transcription factors are essential for growth and development of plants and involved in a plethora of processes. They are a widespread family of transcriptional regulators occurring in multicellular algae, monocots and dicots(40,41). In this study we performed a detailed phylogenetic analysis of barley TCP transcription factors and evaluated the protein-protein interactions of VRS5(HvTB1). One of the identified interactors and closely related protein, HvTB2 showed a similar expression pattern. Targeted mutagenesis showed that HvTB2 is essential for maintaining barley spike architecture. Taken together, this work increases our understanding of the role of TCP transcription factors in shaping barley plant architecture.

TCP transcription factors can form homo- and heterodimers, which affect their DNA binding capacity and specificity. They interact with a plethora of other proteins, including components of the circadian clock and various other transcriptional regulators(40,42). Within the unbiased screen of VRS5(HvTB1) against the Y2H barley cDNA libraries we identified SWItch/Sucrose Non-Fermentable (SWI/SNF) complex subunits. At the protein level, the activity of CIN-like TCPs is known to be regulated by several chromatin remodelling complexes including SWI/SNF(43,44). The interaction of VRS5(HvTB1) with SWI/SNF might point towards a conserved mechanism, where the activity of TB1 is modulated by chromatin remodelling factors at the protein level, similar to the CIN-like TCPs. We also identified other transcriptional regulators such as TCPs and NF-Y amongst the interactors of VRS5(HvTB1) in both the unbiased screen and the heterologous screen against the Arabidopsis TF collection. Large scale Y2H interaction studies in Arabidopsis showed an interaction between AtBRC1 with NFY9(45). NF-Y proteins are a large family of transcriptional regulators known to act in several plant developmental processes and abiotic stress responses(46). It therefore remains to be evaluated how specific the interaction between VRS5(HvTB1) and members of the SWI/SNF chromatin remodelling and NF-Y TF family are. Nevertheless, our analysis shows a glimpse into the putative protein-protein interactome of VRS5(HvTB1)

Further, more detailed analysis using barley class II TCPs show that within the class II TCPs VRS5(HvTB1) preferably heterodimerizes with the CYC/TB1 clade rather than the CIN clade. One of these key putative VRS5(HvTB1) interactors identified is HvTB2 which, similar to VRS5(HvTB1), inhibits the outgrowth tillers. However, some difference in functionality also occurs. While VRS5(HvTB1) inhibits the outgrowth of lateral florets in the main spike through regulation of VRS1(HvHOX1), HvTB2 does not show an obvious row-type phenotype. Instead, HvTB2 suppresses the outgrowth of branches from the main spike. This points towards a mechanism in which VRS5(HvTB1) and HvTB2 are only partially redundant. Taken together, VRS5(HvTB1) heterodimerizes with other transcriptional regulators. To what extent the heterodimerization of VRS5(HvTB1) influences DNA binding and subsequent transcriptional regulation of the target genes remains to be elucidated. Taken together, our analysis opens up the opportunity for expanding the VRS5(HvTB1) interactome and the subsequent identification of other key regulators of plant architecture such as HvTB2.

Genome duplication and diversification has played a major role in the evolution of the TCP transcription factor family. For example, mosses and ferns contain five to six members(41), whereas the dicot model system Arabidopsis has 24(47). The gene duplication events are not always uniform, maize for example mainly shows duplicates in the CYC/TB1 clade. We identified 21 TCP transcription factors in barley and 62 in wheat. Wheat contains mostly three orthologues when compared to barley, representing the hexaploidy nature of the genome. Like in maize and rice, barley and wheat have additional grass-specific duplicates in the TB1/CYC clade. Within this clade both barley and wheat contain close homologues, such as maize *ZmBAD1* and rice *OsTB2* genes, *HvTB2* and *TaTCP24*, respectively. Although these genes are phylogenetically closely related, vast differences in functionality are observed. *ZmBAD1* (also referred to as *WAB1*) is expressed in the pulvinus where it regulates branch angle in the tassel(14,48). *OsTB2* (also referred to as *REP1*) is expressed in the palea primordium during early flower development and in later stages in the stamens and vascular bundles of the lemma and palea(11,12). It is involved in palea development and floral zygomorphy in rice. Recent studies have shown that *OsTB2* is also expressed in the basal tiller node where it induces the outgrowth of tillers(12). *OsTB1* and *OsTB2* act antagonistically on tiller development. *HvTB2* is mostly expressed in the developing inflorescence, where it based on RT-PCR analyses follows a similar expression pattern when compared to *VRS5(HvTB1)*. Targeted mutagenesis of *HvTB2* resulted in spikes that lost their characteristic determinant growth pattern, and exhibited lateral branches arising from the main spike. This suggests that *HvTB2*, in contrast to its rice homologue, acts as branching inhibitor rather than as inducer. In line with this, our phenotypic analysis showed that *hvtb2* mutants exhibited an increased tiller number when compared to the wild type cv. Golden Promise, revealing a more general role as branching inhibitor. In this respect, HvTB2 does not appear to act antagonistically to HvTB1 on tiller development.

Haplotype analysis indicates that *HvTB2* is highly conserved in barley. Considering the phenotype of *hvtb2* it is highly tempting to speculate that *HvTB2* was under selection to maintain spike architecture. The function of *TaTCP24*, which is phylogenetically closely related to HvTB2 and also expressed in developing spikes(49), remains to be elucidated. Taken together, although *BAD1, OsTB2* and *HvTB2* are phylogenetically closely related, they seem to exhibit functional diversity.

RNA *in situ* hybridization shows that *HvTB2* mRNA is localizes at spikelet meristem boundaries. This result, combined with the presence of fused seeds in the generated CRISPR-*hvtb2* knockouts, suggests that HvTB2 plays a role in the specification of the spikelet meristem boundaries. Recently, two independent manuscripts were published while this work was under preparation(50,51). In the first one, published by Shang et al. (2020)(51), the *BDI1* locus was mapped, and the underlying gene corresponded to *HvTB2*. In this work, a significant upregulation of both *SPL8-like* and *HvTB2* was observed in the *vrs4* mutants, while in *hvtb2(bdi1) SPL8* was significantly downregulated at the awn primordium stage. Our RT-PCR analysis also shows that *HvSPL8-like* was significantly downregulated in the *hvtb2* mutants. This suggest that *HvTB2* acts upstream of *SPL8-like*, similar to maize *ZmBAD1(WAB1)* pointing to a conserved mechanism at the molecular level. In a second manuscript, published by Poursarebani *et al* (2020), it was shown that *HvTB2* is the causal gene underlying the *com1* and *int-h* locus, which is corroborated by our analysis. Previously, it was demonstrated that *VRS5(HvTB1)* acts downstream of VRS4(HvRa2), a key regulator of row-type which promotes spikelet and floret determinacy(25,33,52). In maize, RA2 acts upstream of *ZmBAD1*, which is phylogenetically closely related to *HvTB2*. Like *HvTB2, VRS4(HvRa2)* transcript is located in the boundary region(52). Poursarebani *et al*(50), placed *HvTB2* downstream of *VRS4(HvRA2)* and showed a down regulation of *HvTB2* in *vrs4*.*k* at the double ridge and AP/LP stage. This suggests that *VRS4* acts upstream of both *VRS5(HvTB1)* and *HvTB2*, at least in regulating inflorescence architecture. Functional *VRS4(HvRa2)* prevents the outgrowth of lateral florets through activating *VRS5(HvTB1)* and *VRS1(HvHOX1)*, the latter is a well-known conserved inhibitor of lateral floret development(53). As such, *vrs1, vrs4* and *vrs5* single mutants display a six-rowed (*vrs1, vrs4*), or intermediate (*vrs5*), phenotype where lateral florets are developed(5,24,25,52,53). Both *VRS4* and *VRS5* act on lateral floret development trough modulating *VRS1* expression(24,25). In addition to this, *vrs4* mutants show similar to *hvtb2* an outgrowth of lateral branches(25,52). In this respect, it is interesting to note that *hvtb2* did not display an obvious six-rowed phenotype. In line with this, our RT-PCR analysis showed that *VRS1(HvHOX1)* expression was not significantly altered in the *hvtb2* mutants. Taken together, we propose that *VRS4(HvRA2)* acts upstream of *VRS5(HvTB1)* and *HvTB2* in supressing the outgrowth of respectively lateral florets and branches in the main inflorescence.

In conclusion, our analysis and two additional recently published independent studies(50,54) have shown the essential role of HvTB2 in maintaining the characteristic unbranched barley spike.

## Material and Methods

### Phylogenetic analysis

Sequences of Rice, Maize and Arabidopsis TCP TF were downloaded again from the iTAK(26) (http://itak.feilab.net/) and GRASSIUS(27) (www.grassius.org) databases and manually curated for missing TCPs. The barley TCPs were also downloaded from there and checked for missing sequences through a BLAST search against the barley genome using the IPK ViroBLAST (https://webblast.ipk-gatersleben.de/) (55,56). Protein sequences were aligned using MUSCLE (Edgar, 2004) in MEGA version 7. Exome number for barley, wheat and rice were extracted from ENSEMBLE plants. In case of splice variants only one sequence was retained.

To identify homologs of HvTB2, we performed a blastp search using the protein sequence as query in the Phytozome database (https://phytozome.jgi.doe.gov/)(57) against peptide sequences from following species: *Arabidopsis thaliana, Brachypodium distachyon, Carica papaya, Cucumis sativus, Hordeum vulgare, Medicago truncatula, Oryza sativa, Populus trichocarpa, Ricinus communis, Sorghum bicolor, Solanum lycopersicum, Triticum aestivum, Vitis vinifera*, and *Zea mays*. BLAST results were filtered with an E-value cutoff of 1E− 10. The phylogenetic tree of HvTB2 homologues was rooted by using *Selaginella moellendorffii* homolog as an outgroup.

Sequences were aligned using MUSCLE(58) in MEGA version 7(59). A maximum likelihood phylogenetic tree was constructed using RAxML(60), using autoMRE for assessing convergence during bootstrapping. For the phylogenetic tree on all TCPs and the HvTB2 homologues only the convergence test was met after 50 and 100 replicates, respectively. The resulting phylogenetic trees were visualized in EMBL iTOL v4(61).

### Haplotype analysis

Haplotype analysis was performed as described previously(34), using a set of 39 research/breeding lines, 137 landraces and 91 wild barley accessions published in Russell et al. (2016)(39). Further exploration of the natural variation was performed by including the WHEALBI dataset (Whealbi), for which only the SNP matrixes are publicly available(38).

### Construction of the yeast-two-hybrid libraries and protein-protein interaction studies

Barley seedlings, cv. Golden Promise, were grown in controlled greenhouse conditions under long day (LD) conditions (16h, 22°C day; 8h, 18°C night). Samples were taken two hours before the end of the light period to maximize the expression of genes involved in floral organ development and flowering time. The developing seedlings were grown in 96-well trays, and fertilized when necessary. Before sampling the development of the main shoot apex (MSA) was scored according to the quantitative scale by Waddington et al. (1983). This scale is based on the progression of the most advanced floret primordium and carpel of the inflorescence. The reproductive MSA is specified by the appearance of the first floret primordia referred to as the double ridge stage (W1.5-W2.0). Subsequently, the first lemma primordium occur (W3.0) followed by the stamen primordium stage (W3.5), which is characterized by the differentiation of the first floral organ primordia and the stem elongation. The induction of floral organ primordia continues until the awn primordium stage (W5.0). The last stage sampled for the library was W6.0, at this stage the stylar channels are closing. For each stage (W0-W5), at least 10 MSA in three independent biological replicates were pooled. Two Y2H screening libraries were generated one for the early developmental stages (W0-W1.5) and one for the late developmental stages (W2.0-W6). These stages have been selected as VRS5 is mainly expressed in the developing shoot apex. All MSA harvested for RNA extraction were frozen immediately in liquid nitrogen and stored at − 80°C. RNA was isolated as described previously(33). Libraries were constructed using the CloneMiner^™^II kit, according to manufacturer’s protocol One exception was the propagation of the libraries in E.coli, which was done on large 150 mm in diameter petri dishes instead of liquid medium. The pDEST22 vector was used as prey vector, and thus the destination vector for the libraries. The resulting libraries contained a titer of 8.87 x 10^6^ and 1.73 x 106 CFU. The variation of the genes in these libraries has been tested by colony PCR followed by sequencing of the PCR amplicon.

Primers targeting the TCP transcription factors used in the yeast-two hybrid screen are listed in Supplemental Table S6. The corresponding TCPs were amplified using Q5® High-Fidelity DNA Polymerase (New England Biolabas) from the cDNA screening library and cloned into pDONR201. For VRS5(HvTB1) the HvTB1-a and HvTB1-b alleles were amplified from cDNA of respectively, cv. Morex and cv. Bowman. Subsequently, the TCP TF were cloned into the bait (pDEST32) and prey (pDEST22) vectors. To prevent auto activation in the bait constructs (pDEST32) the N-terminal part of the full length TB1 protein was removed (VRS5(HvTB1^NtDEL83^)). The TCP domain was kept intact for all construct used. Autoactivation was tested on selective medium containing -L + 3AT and -LA. Only the -LA marker showed no autoactivation (S4A Figure), and therefore the screen was performed using -LWA medium. As negative control, HORVU.MOREX.r2.3HG0240550 was included, which is annotated as a transcriptional regulator without known domains. As positive control, all plates were grown on media containing -LW in parallel to the selective -LWA plates. For the heterologous screen against the Arabidopsis TF library (32), VRS5(HvTB1^NtDEL83^) was used as bait and the library as prey. Subsequently, the screen was performed on -LWA medium using medium containing -LW as positive control, as described above.

### CRISPR-CAS mutagenesis

For CRISPR-CAS mutagenesis OsU3 promoter, which used a “A” as start site, was used, which was linked in the Golden Gate vector system(62). In total three guides were used. guide 1: GCAGCTTCTCCATGGCGCCT; guide 2: GCTCCTCCTCTGGCGGACAT; guide 3: ACTGGCGCAGTGCAGGCCGC. Plants were transformed as described previously(63). The resulting primary transformants were selected for presence of CAS9. In the second generation, two lines were selected based on mutational events. Transformants were genotyped using the Phire Plant Direct PCR Kit (Thermo Fisher Scientific), amplified fragments were directly sequenced. Primers for genotyping the generated mutants are added in Supplemental Table S6.

### Plant growth and phenotyping

For plant phenotyping between cv Golden Promise (GP) and *hvtb2* mutants plants were grown on soil at 22°C during the day (light, 16 hours) and 16°C during the night (in darkness, 8 hours) in 1 L pots supplied with fertilizer and water when needed. Tiller number was recorded at full maturity. Thousand grain weight (TGW), grain number per spike and size were recorded after drying of the spike/seeds. Statistical analyses were performed using the statistical software R (http://www.r-project.org/) Differences between wild type and mutant genotypes were determined using a student’s t-test or a one-way ANOVA combined with a Tukey HSD for multiple comparison

### RNA *in-situ* hybridization, EM microscopy and RT-PCR analysis

Plants were grown on soil at 22°C during the day (light, 16 hours) and 16°C during the night (in darkness, 8 hours) in small 40-well trays. Probes for HvTB2 mRNA were prepared from the whole coding sequence (start to stop codon). Cloning and RNA probe synthesis was performed as described Kirschner et al. (2017)(64) and used as full-length RNA probes or with a subsequent hydrolysation to 150 bp. RNA in situ hybridizations on shoot apical meristems of the double ridge stage (and the awn primordium stage were performed as described before(64).

For scanning electron microscopy, dissected main inflorescences were mounted on a copper specimen holder with freeze hardening glue and frozen in liquid nitrogen. Images were obtained using a FEI Magellan 400 microscope, which is equipped with a Leica cold stage for cryo-microscopy. Low-temperature SEM was performed on the frozen shoot apical meristems. Images of *hvtb2-1* and *hvtb2-2* were processed using Adobe Photoshop to colour code the outgrowing side shoots. Staging of the apex over development was done using a standard binocular microscope. For RT-PCR and monitoring the shoot apex development, plants were grown in 96-well trays, under controlled greenhouse conditions as described above. Leaf-enriched developing inflorescences were collected lemma and stamen and awn primordium stages. RNA-isolation for RT-PCR analysis was done using the PureYield^™^ RNA kit (Promega). For expression analysis of HvTB2 in *vrs4* background the *vrs4*.*k* mutant (GSHO 1986), which is a near isogenic line in cv. Bowman. All RT-PCR experiments were done in at least three biological replicates. Statistical differences were calculated using a two-tailed unpaired Student’s t test. Primers for RT-PCR analysis are included in Supplemental Table S6.

## Acknowledgments

We cordially thank: Rients Niks for sharing greenhouse space; Marcel Giesbers, for generating the EM images of the barley apex; Agatha Walla for assistance in the haplotype analysis; Steven Groot for making pictures wild type and mutant seeds, awns and spikes; and Martijn Wiekens for technical assistance in generating the Y2H libraries during his BSc thesis work; and Peter van Esse and Mark Youles for sharing vectors used for CRISPR-CAS mutagenesis. This work has been financially supported by the Gatsby Foundation (MJM) and VENI (WvE, project number 15060).

## Supporting information captions

**S1 Fig. Genomic organization of the TCP TF gene family in barley**. Red boxes represent coding exons, white boxes exons and lines represent introns. Wheat and rice TCPs that have a different genomic organization when compared to barley are highlighted in blue.

**S2 Fig**. Maximum likelihood phylogenetic tree of *HvTB2-like* genes in 19 monocot and eudicot plant species. Tree was build using the protein sequences. The sequence of a TCP homolog obtained from Selaginella moellendorffii (transcript ID 89227) was used for rooting. The barley *HvTB2* gene described in this study, HvTB2, is highlighted in red. Functionally characterized genes within the same clade are marked in blue. Arabidopsis class II TCPs in the eudicot branched are marked in green. Bootstrap support (%) is shown at the nodes. Abbreviated species names are given before gene identifiers. Aet: *Aegilops tauschii*; Ath: *Arabidopsis thaliana*; Bd: *Brachypodium distachyon*; Cp: *Carica papaya*; Cs: *Cucumis sativus*; Hv: *Hordeum vulgare*; Mt:*Medicago truncatula*; Os: *Oryza sativa*; Pt: *Populus trichocarpa*; Pv: *Phaseolus vulgaris*; Rc: *Ricinus communis*; Sb: *Sorghum bicolor*; Sc: *Secale cereale*; Si: *Setaria italica*; Sl: *Solanum lycopersicum*; Ta: *Triticum aestivum*; Vv: *Vitis vinifera*; Zm: *Zea mays*. Scale bar = 0.1 substitutions per site.

**S3 Fig. Barley apex and crown tissue used isolated to generate yeast-two-hybrid libraries**. Main shoot apex (MSA) and crown tissue of developing barley seedlings was excised at different developmental stages. Library 1 was made from crown tissue including the vegetative apical meristem. Library 2 was made from shoot apical meristem tissue obtained during various stages of floral organ development, staring at the floral transition which is marked by the double ridge. The last samples for library 2 were taken after the induction of floral organ primordia was completed. Random PCR amplification of the inserts present in several independent colonies. indicated that the libraries include cDNA fragments between 500-2000 bp. Sequencing of 10 colonies verified that there was a good variation in the identified proteins.

**S4 Fig. Detailed overview of yeast-two-hybrid protein-protein interactions. (**A) Table showing the results of the autoactivation test. For each construct at least three colonies were scored. (B) Number of replicates performed in the protein-protein interaction studies (top panel), compared to the number of interactions observed (middle and bottom panel). Each interaction was scored in at least 6 independent replicates. Differences between replicates are visualized by dividing the interactions scored by the number of replicates with: a score of 0 (yellow) no interaction; and a score of 1 (dark green) always an interaction; values in-between 0-1 indicate the constancy of the results between replicates.

**S5 Fig.Target region for CRISPR-CAS mutagenesis of *HvTB2***. Black bars mark the three guides, red triangles the NGG PAM recognition site. The orange block shows the TCP domain.

**S6 Fig.Quantification of the *hvtb2* spike phenotype**. Seeds per rachis internode on each side of the spike are indicated in green. GP did not contain any lateral spikelets (gray) nor lateral branches while in *hvtb2* most of the basal lateral spikelets are developed into seeds (pink). Purple bloc ks indicate branches occurring at the rachis node. C indicates central spikelet, L indicate lateral spikelets.

**S7 Fig. Phenotypical analysis of the *hbtb2* mutant. (**A) Original images used to generate the subpanel 3B. Three representative seeds with awn were selected to visualize the difference in awn architecture for seeds on the basal part of *hvtb2* mutants when compared to the wildtype cv. Golden Promise (GP). (B) Phenotype of *hvtb2* compared to GP, seeds are removed for better visualization of the branches. (C) Number of grains per spike, n= 9 spikes; D-F) Seed parameters TGW (n=20); grain width (n=30; and grain length (n=30). In the *hvtb2* mutants the central and lateral seeds were measured separately. Statistical differences are based on a one-way ANOVA, combined with a combined with a Tukey HSD for multiple comparison. Letters indicate differences when compared to GP using a; P ≤ 0.05.

**S8 Fig. Genotyping *int-h* and *com1*. (**A) Table showing the PCR amplification results of *int-h* and *com1* lines. Green marks regions where a PCR amplicon was obtained, gray indicates no amplicon. Top panel shows the regions that were targeted in the PCR analysis: promoter region of *HvTB2* (dark green), coding sequence (blue) and downstream region (orange). Up to a region of 2,500 base pairs upstream of the start and 7050 bp downstream of the start no PCR amplicon was obtained in *int-h*.*42, int-h*.*43* and *int-h*.*44* as well as com1.a and *com1*.*b*. Primers used in the assay are included in Supplemental Table S6. (B) int-*h*.*83* and *com1*.*c* contained a non-synonymous polymorphism within the conserved TCP domain (black box) which was not present in the wild type control nor identified as common haplotype. (C) PROVEAN score, which predicts whether an amino acid substitution or indel has an impact on the biological function of a protein indicates that there is no effect of the observed haplotypes HAP2, 7 and 8 while the SNPs in *int-h83* and *com1*.*c* are predicted to have deleterious effects.

**S9 Fig. Shoot apical meristem development of hbtb2-1 compared to cv. Golden Promise. (**A) development of the shoot apex of wildtype and *hvtb2-1* mutant. At double ridge stage no differences were observed while at awn primordium stage a clear outgrowth of the lateral branch is observed. (B) Shoot apical meristem development of cv Golden Promise (GP) versus *hvtb2-1*, monitored using the Waddington scale (W). (C) leaf number of *hvtb2-1* compared to the wildtype GP. For both (B) and (C) n≥6 plants. No significant differences were observed.

**S10 Fig. In-situ hybridization in cvBowman targeting HvTB2**. The RNA *in-situ* hybridization was performed at the double ridge stage, (A) and the awn primordium stage (B). The first two images in panel B show the original compiled images used for figure 4B, whereas the third image shows the same tissue but a different sectioning depth.

**S11 Fig. Haplotype analysis of *HvTB1*. (**A) Haplotype network of *VRS5(HvTB1)* comprising elite, landrace and wild barley lines. In total sequences of 670 different genotypes were included in the analysis. The two major haplotypes observed HAP1 and HAP2 give a clear distinct between two-rowed and 6-rowed architecture signifying the selective advantage of these haplotypes for specific backgrounds. (B) Representation of the two main VRS5(HvTB1) haplotypes corresponding to the 2-rowed and 6-rowed cultivars. (C) PROVEAN score, which predicts whether an amino acid substitution or indel has an impact on the biological function of a protein indicates that there is no effect of the observed haplotypes.

**S1 Table** Genomic location of TCP transcription factors in barley

**S2 Table** Gene names and identifiers used to construct the TCP phylogenetic tree (Figure 1)

**S3 Table** Screen of HvTB1 protein against a Y2H screening library in yeast **S4 Table** Expression of TCP transcription factors used in the Y2H screen in the developing shoot apex

**S5 Table** Measured phenotypical data for wildtype Golden Promise, hvtb2-1 and hvtb2-2

**S6 Table** Primers used in this study

